# Oncogenic Phase Transitions: How Mutant p53 Drives Amyloid Formation in p63 and p73 Liquid Droplets

**DOI:** 10.1101/2024.03.28.583974

**Authors:** Elaine C. Petronilho, Guilherme C. de Andrade, Gileno dos S. de Sousa, Fernando P. Almeida, Michelle F. Mota, Mayra A. Marques, Tuane C. R. G. Vieira, Guilherme A. P. de Oliveira, Jerson L. Silva

**Author notes:** Equally first authors (Elaine C. Petronilho, Guilherme C. de Andrade).

## Abstract

Phase separation (PS) of p53 is critical in its path to amyloid aggregation, a process linked to cancer. P63 and p73 exhibit dual roles as tumor suppressors and oncogenes and are often heightened in tumors. Their coaggregation has been proposed in cancer, yet their PS contribution remains unknown. This study investigates the phase behaviors of p53, p63, and p73. P63 and p73 undergo liquid-liquid phase separation (LLPS). Unlike p53, p63 and p73 do not form amyloids under increased temperatures, underscoring an aspect of the processes involved in cancer. Unlike p63 and p73, wild-type and the M237I mutant p53 initially form droplets at 4°C, but at temperatures up to 37°C, they begin to aggregate and bind to Congo red, showing amyloidogenesis. Intriguingly, mutant p53 promotes amyloid-like states in p63 and p73 and hijacks p73 into membrane less organelles. Wild-type p53 has a moderate effect on p63 and p73’s amyloid aggregation. Additionally, heparin prevents the prion-like aggregation of p63 and p73 induced by p53. Our results shed light on how mutant and wild-type p53 may trigger amyloid aggregation of p63 and p73, revealing the capacity of p53 amyloid droplets within cancer. These insights expand the possibilities for developing cancer therapies targeting the prion-like conversion of p63 and p73 influenced by mutated p53.

**Relevance:** The prion-like action of mutant p53 on p63 and p73 droplets would be the basis for an oncogenic gain of function of the p53 mutation.

## Introduction

The p53 gene, often dubbed the “guardian of the genome,” has long been recognized as having a central role in cancer biology^1, 2^. With mutations in this pivotal tumor suppressor occurring in over half of all malignancies, it is crucial to search for innovative treatments targeting these anomalies^3^. It has been found that mutant p53 tends to undergo amyloid aggregation, which is correlated with the carcinogenic process^4–7^. Our group and others have contributed to this growing body of knowledge, uncovering the threatening ability of mutant p53 to confer oncogenic properties, including chemotherapy resistance, upon these aggregates^4, 8–11^. Protein aggregation and misfolding cause a toxic gain of function in neurodegenerative diseases, while in cancer, mutant aggregates of p53 result in a loss of its tumor-suppressing function and enhance cells’ oncogenic potential^6, 12–15^. This revelation has transformed mutant p53 into a novel pharmacological target, one that had previously been considered elusive^11, 12, 16^. This pioneering research, established in different laboratories, laid the groundwork for understanding the pathological role of mutant p53 amyloid aggregates, observed in breast cancer cells and other tumoral diseases, including skin, prostate, and ovarian carcinomas^17–20^.

Biophysical studies have revealed that despite the structural similarities between the DNA binding domains of p53 and its paralogous forms p63 and p73, the latter two exhibit a reduced propensity for aggregation^21, 22^. Biophysical studies and molecular dynamics (MD) simulations revealed specific regions of increased exposure backbone hydrogen bonds (BHBs) for p53 compared with p63 and p73^21^. The regions of structural vulnerability in the p53 DBD are new targetable sites for modulating p53 stability and aggregation, a potential approach to cancer therapy^11, 21^. This difference pinpoints a unique aspect of p53’s structure that may be exploited therapeutically. Thus, the modulation of p53’s stability and aggregation present a tempting target for cancer therapy, with recent studies showing how mutant forms of p53 can interact with p63 and p73 through a co-aggregation mechanism^7, 23^. Mounting evidence confirms that p53 mutants associate with p63 and p73, impairing the tumor suppressor functions and increasing oncogenicity^7, 11, 24^. However, the mechanisms of the interaction and oncogenic conversion are not fully understood.

The phenomenon of phase separation has recently gained prominence as a crucial event in the life cycle of a cell, with implications for oncogenicity^11, 25–31^. We have recently demonstrated that phase separation is a precursor to the amyloid aggregation of p53, presenting a significant avenue for therapeutic intervention^31^. The tendency to undergo phase separation and, eventually, phase transition seems to be an intrinsic property of p53^10, 29, 31–34^. Other p53-related proteins and tumor suppressors equally undergo phase separation in the nucleus^35, 36^. In the case of the tumor suppressor speckle-type BTB/POZ protein (SPOP), mutations have been associated with phase separation defects^35^. SPOP mutations disturb substrate interactions that affect SPOP phase separation and its localization to membranelles organelles. p53 has arisen as a cancer-related protein in which mutations can disturb its folding, likely leading to phase transition and, thereby, to the formation of amyloid aggregates with oncogenic properties (GoF)^11, 31, 37^.

When unraveling the intricate protein interactions within cancer, the interplay among p53, p63, and p73 emerges as a central focus of our study. Our research endeavors to unravel the phase behaviors of these proteins, illuminating the complex mechanisms that drive their transition from a liquid phase to an amyloid-like solid state. Our findings contribute to understanding how the amyloid aggregation of p63 and p73 is precipitated by both the mutant M237I and wild-type forms of p53. Such insights offer new vantage points regarding mutant p53’s pathogenic influence in oncogenesis, setting the stage for groundbreaking therapeutic strategies. The induction of amyloid-like conformation by p53 variants in paralogous proteins holds significant cancer-causing potential. Heparin, a polyanion known for its regulatory influence on p53 aggregation, emerges as a potent inhibitor of the prion-like aggregation of p63 and p73 mediated by p53. Transfected cells with M237I mutant evidence the colocalization of p53 condensates with p73. By pinpointing the specific conditions conducive to the amyloid aggregation of p63 and p73, our work enhances the comprehension of p53’s role in cancer development and underscores the therapeutic potential of intervening in these phase transitions as a novel anticancer strategy.

## Results

We initiated our investigation by evaluating the phase separation behavior of critical proteins implicated in cancer, namely p63, p73, and p53. Specifically, we examined the propensity of the DNA domains of these proteins, denoted as p63C, p73C, and p53C, respectively, to undergo liquid-liquid phase separation across a range of protein concentrations. As illustrated in Figure 1a, both p63C and p73C, along with p53C, exhibited liquid-liquid phase separation at various protein concentrations. This process was facilitated by polyethylene glycol (PEG) at a temperature of 4°C and was observed using differential interference contrast (DIC) microscopy (Figure 1a-f and Supplementary figure 1). Visual inspection of the images showed that demixing of p63C occurred at lower concentrations than p73C and showed the tendency to form larger droplets in increased concentrations (Figure 1e). Expanding upon our prior research findings, which elucidated the phase separation of p53C leading to aggregation across diverse conditions^31^, we observed that the M237I mutant phase separates in the presence of PEG and transitions to a solid-like state more rapidly than the wild-type p53C (Figure 1c and f). Within living cells, mutant full-length p53 transfection resulted in phase separation and phase transition within nuclear compartments, likely contributing to gain-of-function effects^31^. Figures 1c and f illustrate that in the presence of PEG, p63C and p73C also formed liquid droplets akin to p53C. After increasing the temperature to 37°C, wild-type and mutant p53C formed aggregates (Figure 1c, arrows). Notably, wild-type and mutant p53C transited from droplets to aggregates (Figure 1f, asterisks), whereas p63C and p73C did not demonstrate intrinsic aggregation (Figure 1c and f). Figure 1d presents the transition of p53C to a solid state at 25°C, capturing the formation of p53C aggregates through successive bright-field DIC microscopy images and video recordings (Supplementary figure 2), demonstrating the seeding mechanism in aggregate formation.

**Figure 1.**
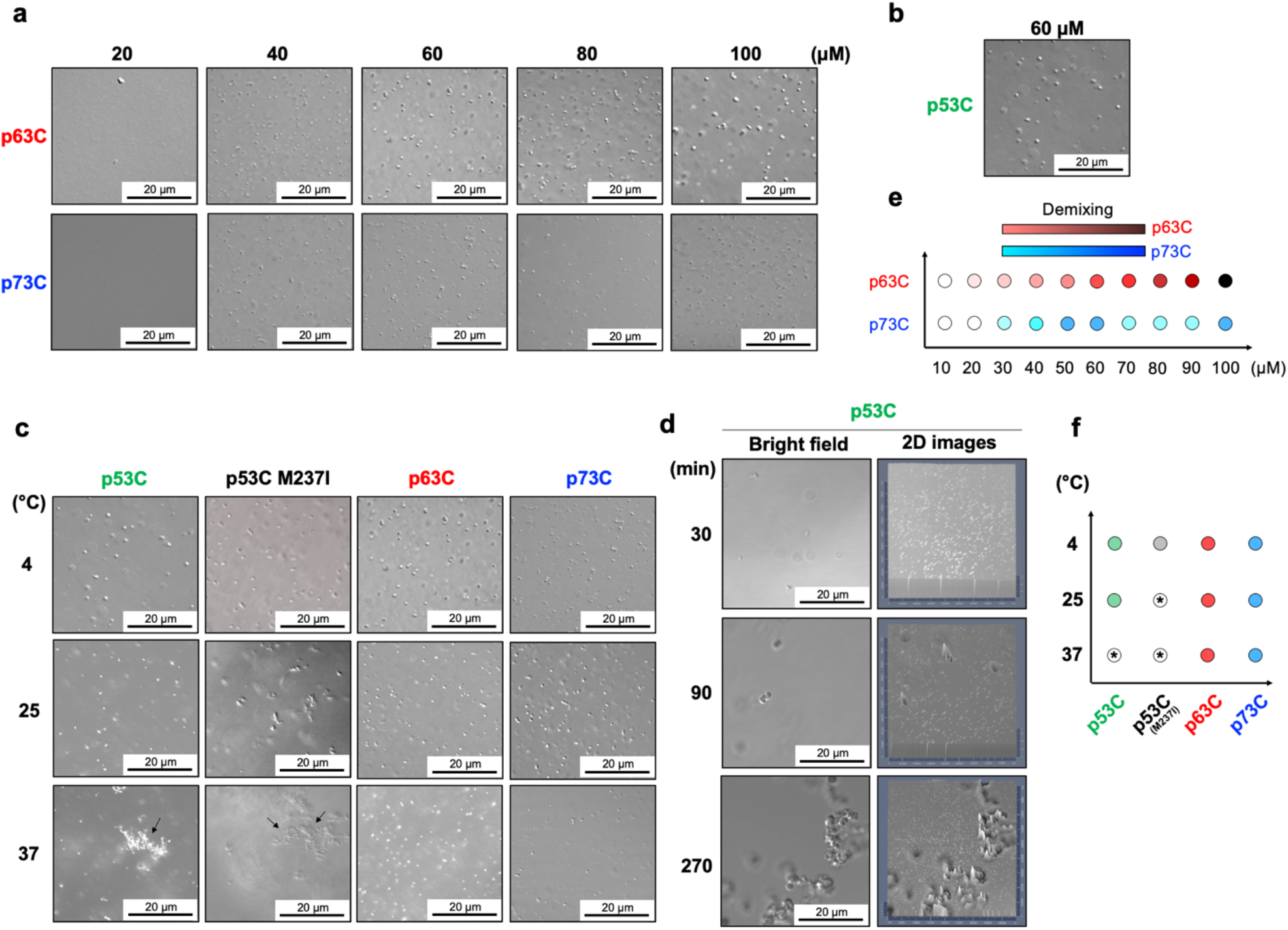
The p53 family phase behavior at the microscale resolution as a function of protein concentration and temperature. Bright-field differential interference contrast (DIC) microscopy images were collected, revealing (a) the concentration-dependent demixing of p63C and p73C, and (b) the demixing of p53C from the PEG solution. (c) The temperature dependence of p53C, M237I, p63C, and p73C demixing and aggregation (arrows). (d) The time dependence of p53C aggregation from droplets at 25 °C (see methods); (e, f) Schematic representations of (e) p63C and p73C demixing concentrations, and (f) the temperature dependence of the studied proteins. Colored circles indicate the phase separation of p53C (green), M237I (gray), p63C (red), and p73C (blue). Circles filled with an asterisk depict aggregation conditions.

We analyzed Congo Red (CR) binding to p53 family proteins at both 4°C and 37°C, which serves as a hallmark of amyloid aggregates. The amyloid nature of p53C aggregates, whether mutant or wild type, was readily indicated by CR binding (Figure 2a, b), a phenomenon not observed for p63C and p73C (Figure 2c, d). Control experiments without PEG revealed no droplets and aggregation at 4°C for p53C and M237I. At 37°C, CR-positive aggregates were observed for p53C and M237I, but not for p63C and p73C (Supplementary figure 3). Light scattering (LS) assays were employed to quantitatively compare the aggregation of p53C droplets versus the non-aggregation of p63C and p73C (Figure 2e-h). The results underscore that p63C and p73C displayed no temperature-dependent aggregation, whether in solution or the liquid droplet state (Figure 2g, h), a striking disparity compared to p53C and M237I. Conversely, p53C and M237I demonstrated temperature-dependent aggregation, evident in both the presence and absence of PEG (Figure 2e, f). Turbidimetry assays at 600 nm further corroborated the results obtained from light scattering analysis (Figure 2i-l) and ThT binding analysis (Figure 2m-p). The observed higher propensity of p53 for temperature-induced aggregation compared to p63C and p73C indicates significant differences in their aggregation behaviors, potentially bearing critical biological implications.

**Figure 2.**
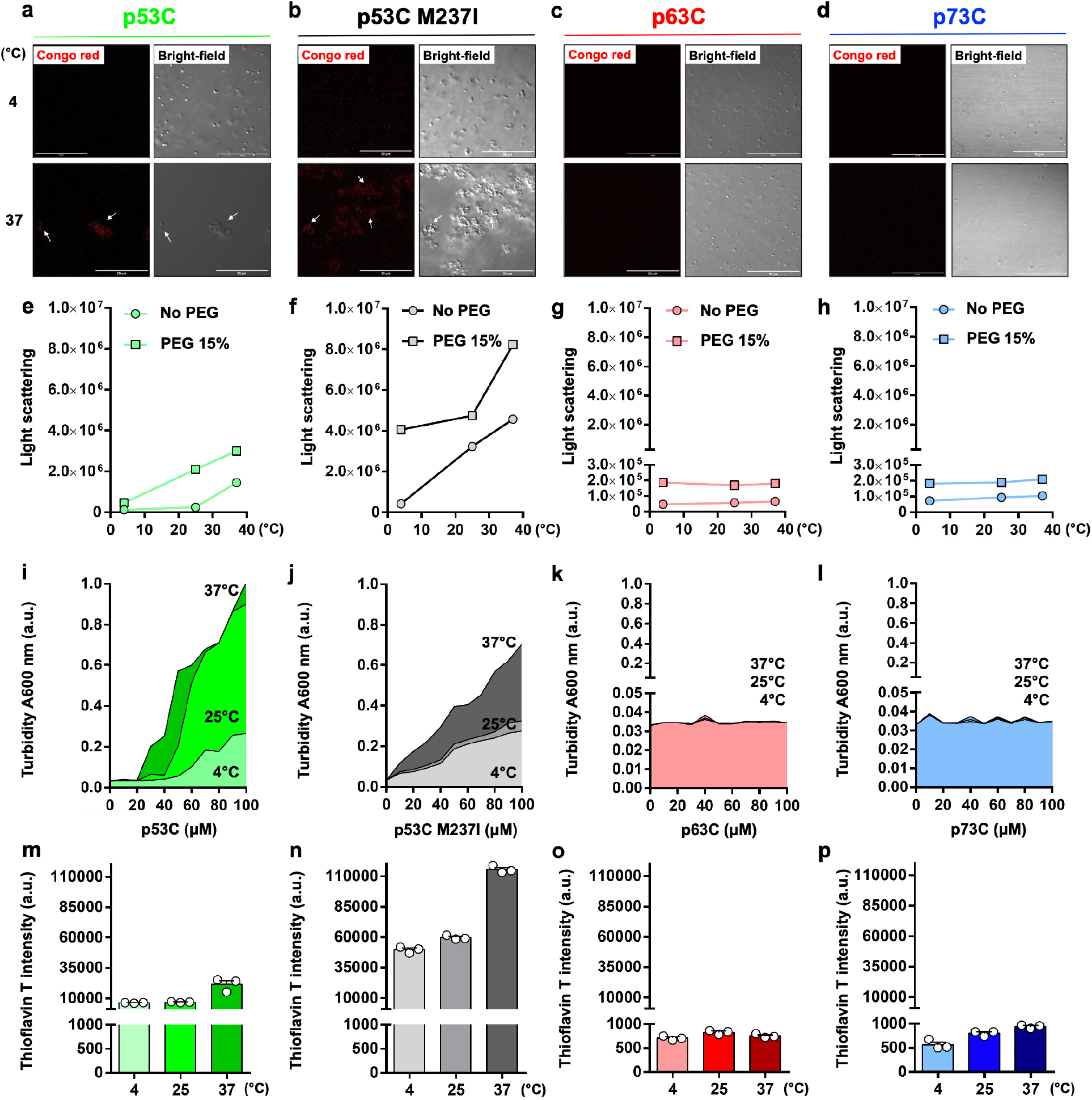
The amyloid nature within the p53 family phase separation and transition. (a-d) Microscopy images were captured, presenting the Congo red (CR) fluorescence and bright-field channels of (a) p53C, (b) M237I, (c) p63C, and (d) p73C at 4 and 37°C. White arrows indicate the presence of p53C and M237I aggregates. Scale bar: 20 μm; (e-h) Line plots showing the area under the light scattering curve as a function of increasing temperatures (4, 25, and 37°C) of (e) p53C (green), (f) M237I (gray), (g) p63C (light red), and (h) p73C (light blue) in the absence (colored circles) and the presence (colored squares) of PEG. Plots in (g) and (h) show two segments in the y-axis to highlight negligible scattering. (i-l) Line plots with the filled area showing turbidity at 600 nm as a function of increasing concentrations of (i) p53C (green palette), (j) M237I (gray palette), (k) p63C (red palette), and (l) p73C (blue palette) at 4, 25, and 37°C. Plots in (k) and (l) show two segments in the y-axis to highlight negligible turbidity. Red and blue palettes are not depicted in (k) and (l) due to similar values across studied temperatures. (m-p) Scatter dot plots showing the thioflavin T fluorescence as a function of increasing temperatures (4, 25, and 37°C) of (m) p53C, (n) M237I, (o) p63C, and (p) p73C. The color palette is the same as in (i-l). Plots show two segments in the y-axis to evidence large changes in the thioflavin T signal across studied proteins. The data are shown as the mean ± s.e.m. of n = 3 independent experiments using the same protein batch.

Figure 3 explores the capability of p63C and p73C droplets to transition into amyloid aggregates in the presence of p53C aggregates, utilizing rhodamine-labeled p63C (Figure 3a) or p73C (Figure 3b). Microscopic examination unveiled that at 37°C, p53C induced the aggregation of both p63C and p73C. Rhodamine-labeled p63C or p73C aggregates colocalized with Alexa 488-labeled p53C (both wild-type and M237I mutant) (Figure 4a-d), suggesting a prion-like seeding process, where mutant p53 may induce coaggregation with typically non-amyloid p63C and p73C. These results provide insight into the gain-of-function phenomena observed in mutant p53 tumors. Figures 4e and 6 summarize the coaggregation dynamics of p63C and p73C in the presence of p53C and M237I.

**Figure 3:**
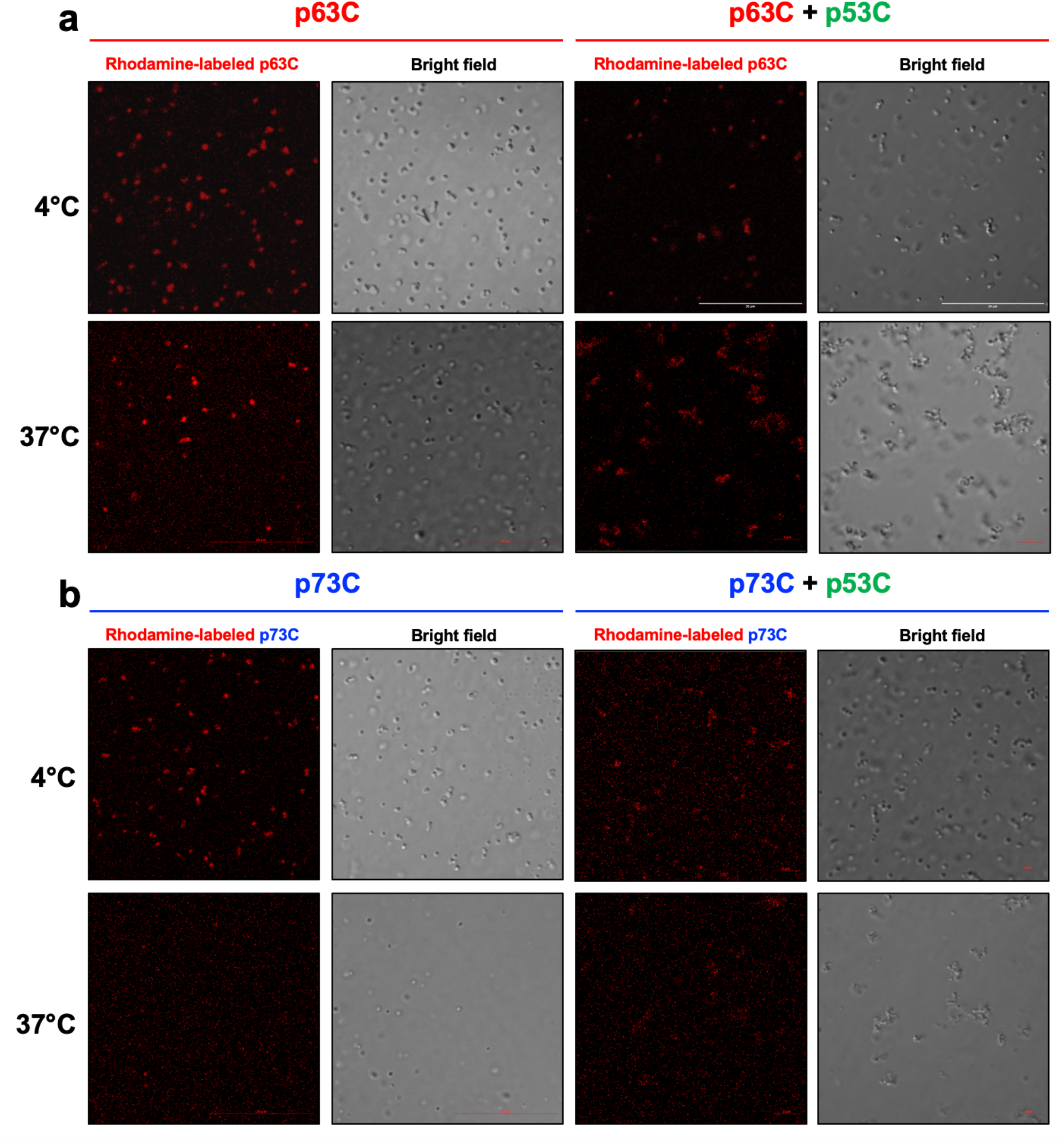
Influence of p53C on p63C and p73C aggregation. Microscopy images showing the rhodamine-labeled (a) p63C and (b) p73C fluorescence and bright-field channels before (left panel) and after (right panel) mixing with nonlabelled p53C at 4 and 37°C. Scale bar: 20 μm.

**Figure 4.**
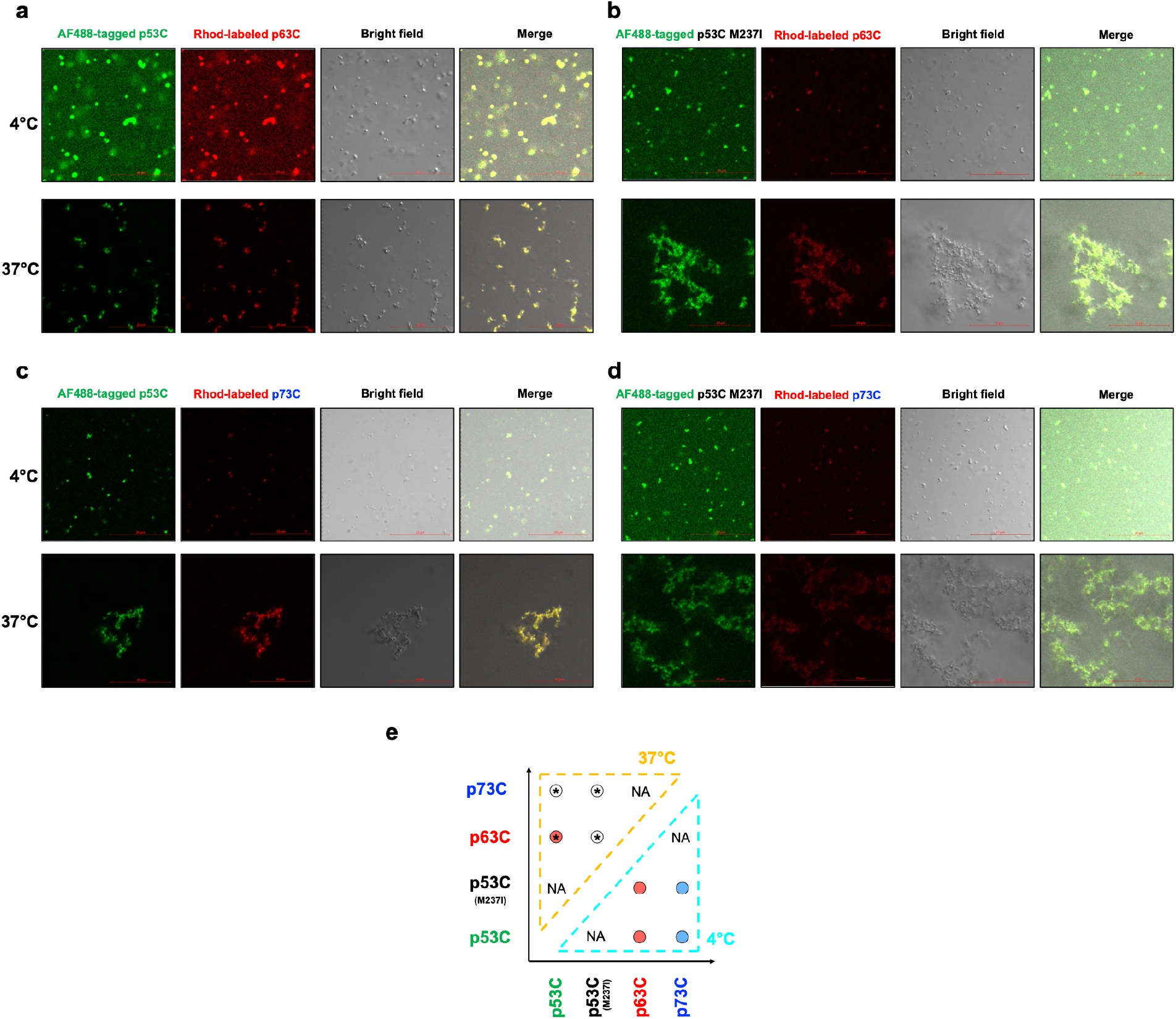
Aggregation of p63C and p73C induced by p53C and M237I. (a-d) Microscopy images showing the Alexa-fluor 488 (AF488) labeled (a, c) p53C or (b, d) M237I (green), the rhodamine-labeled (Rhod) (a, b) p63C or (c, d) p73C (red) fluorescent channels together with the bright-field and merged images at 4 and 37°C. Scale bar: 20 μm. (e) Schematic representation of the crosstalk within the p53 family members at 4 (dashed cyan) and 37°C (dashed orange). Red and blue circles depict demixing with droplet formation. Red and white circles filled with an asterisk indicate a mixture of droplets and aggregates, and mainly aggregates, respectively. NA stands for not analyzed.

Light scattering measurements also evaluated the prion-like seeded aggregation of p53C (Figure 5a-c). P53C (wild-type and M237I mutant) was initially incubated at 37°C, and then the aggregated protein was diluted fortyfold to be used as a seed and mixed with p63C. P53C seeds increased p63C light scattering in the presence of PEG (Figure 5a-c). The M237I p53C mutant exhibited a higher seeding capacity than the wild type (Figure 5c). The results, particularly with the M237I mutant, demonstrated that p53C seeds can trigger p63 aggregation even in significantly reduced concentrations.

**Figure 5.**
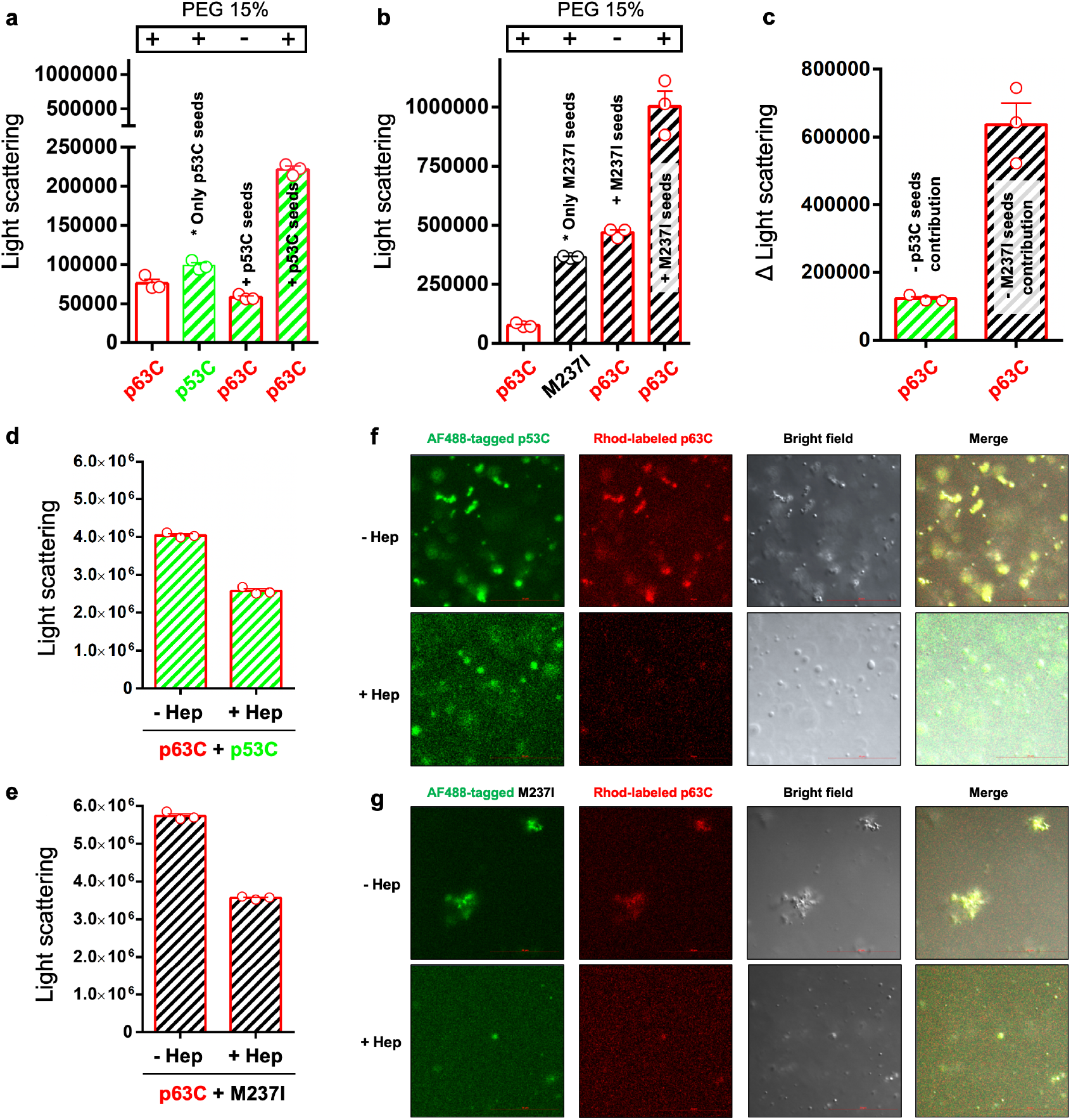
p53C seeds p63C droplets conversion to aggregates and heparin inhibition. (a-e), Scatter dot plots showing the area under light scattering (LS) curves of (a, b) p63C (red bars), seeds of (a) p53C (green bar) or (b) p53C M237I (black bar), after mixing p63C with (a) p53C seeds (red bars colored green) or (b) p53C M237I seeds (red bars colored black), in the absence (-) or presence (+) of PEG 15%. (c) Changes in the LS (Δ area) show the p63C aggregation in the presence of PEG after subtracting the p53C (red bar colored green) or M237I (red bar colored black) seeds contribution. (d, e) the effect of heparin on p63C aggregation when mixing p63C with (d) p53C or (e) p53C M237I in the presence of PEG. The data are shown as the mean ± s.e.m. of n = 3 independent experiments using the same protein batch. (f, g) Microscopy images showing the AF488-labeled (f) p53C or (g) M237I, the Rhodamine-labeled p63C, the bright-field, and merged channels in the absence (-) or presence (+) of heparin. Scale bar: 20 μm.

To elucidate the colocalization of the M237I p53 mutant with p73, we employed a specialized transfection protocol on p53-null H1299 carcinoma cells. Previous studies have shown that the full-length M237I p53 mutant tends to undergo phase separation in the nucleolus, leading to the formation of gel-like droplets^31^. Figure 6 illustrates that the transfected cells displayed p73 puncta colocalizing with EGFP-tagged p53 M237I within the nucleolar membrane-less compartments (see also Supplementary Figure 5). It is noted that colocalization of p73 and the p53 M237I mutant was less common in other nuclear areas.

**Figure 6.**
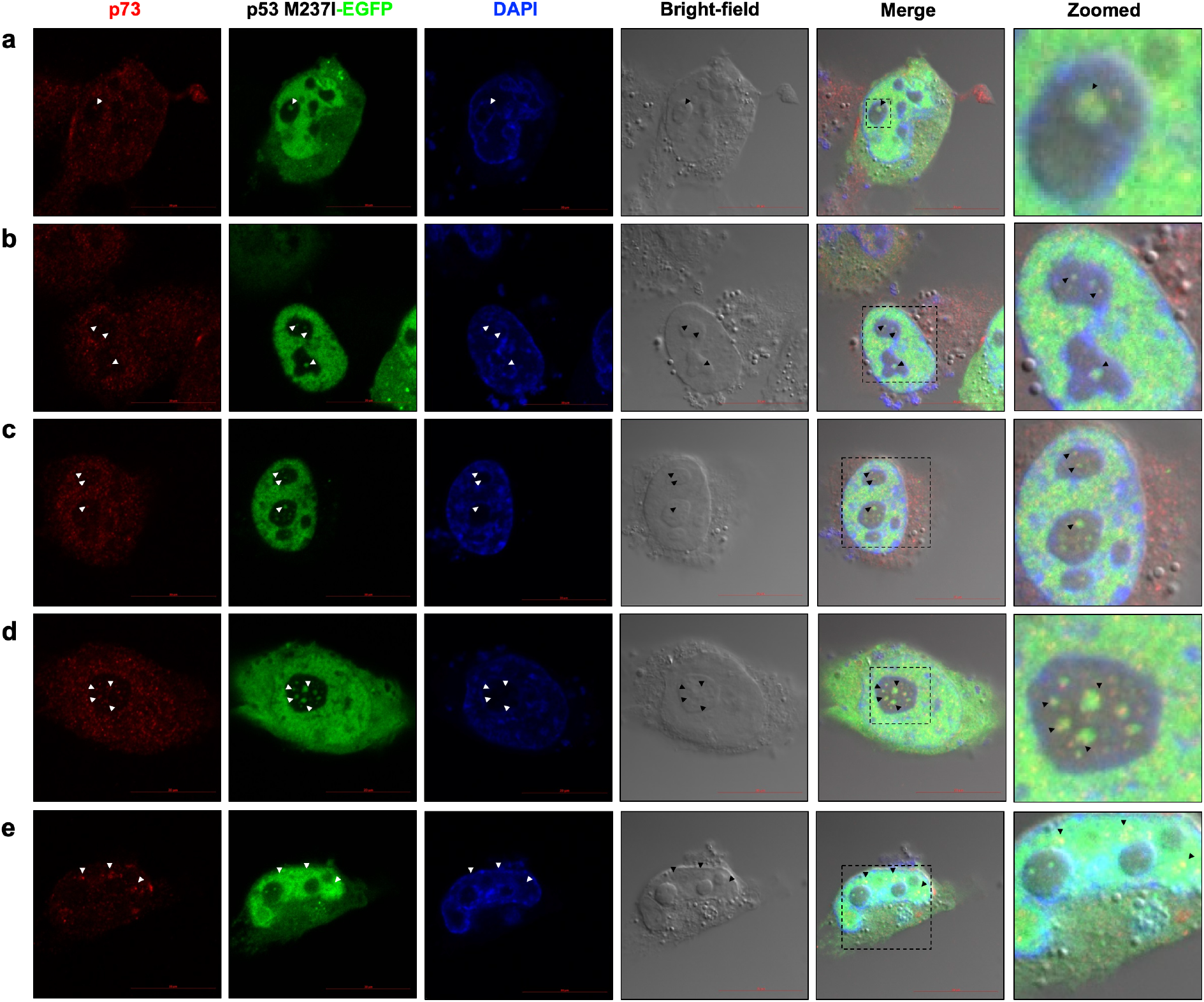
Collection of microscopy images showing (a-e) five distinct regions of interest and the corresponding channels: p73 (red), EGFP-tagged p53 M237I (green), nuclei (blue), bright-field, and merged. Last panel show zoomed images from dashed rectangles to emphasize p53 and p73 colocalization spots (white and black arrowheads).

Collectively, the coaggregation phenomena depicted in Figures 3 and 4 and the seeding data from Figure 5 highlight a prion-like mechanism. This underscores the capability of mutant p53C to facilitate pathological amyloid aggregation in p63C and p73C proteins, which generally do not form such aggregates. This coaggregation activity, potentially contributing to the oncogenic gain-of-function observed in tumors harboring mutant p53, was revealed through our detailed microscopic analysis. These findings illuminate the potential for mutant p53 to modulate the phase behavior of p63 and p73, implicating it in the pathological aggregation processes tied to cancer.

The comprehensive insights derived from Figures 3 to 5 contribute to an enhanced understanding of the amyloidogenic propensities of p53 mutants, suggesting potential avenues for novel therapeutic interventions in cancer. One such approach to disrupt the amyloidogenic process involves the utilization of inhibitors targeting p53 aggregation. Our previous research found that heparin modulates p53 phase separation, inhibiting its progression to a more solid state and likely maintaining it in a dynamic, gel-like state^31^. Consequently, we explored whether heparin could similarly influence the prion-like seeding property of p53C (Figure 5d-g). The data revealed that in the presence of heparin, M237I p53C seeds fail to induce the transition of p63 and p73 from droplets into aggregates (Figure 5f, g and Supplementary Figure 4). Light scattering measurements provided further support, demonstrating that both wild-type and mutant p53C diminished p63C aggregation in the presence of heparin while likely leaving the p63C droplets unaffected (Figure 5d-g). These findings underscore the potential of heparin as an inhibitor of p53-mediated aggregation of p63C, highlighting its promise as a therapeutic intervention in cancer.

## Discussion

Our results feature the fundamental role of phase separation in the pathophysiology of p53-related cancers. The transition of p53 from a liquid to a solid state, observed in both mutant and wild-type forms, corroborates the hypothesis that protein phase behavior is intrinsically linked to cellular dysfunction in cancer^11^. The novel finding that p53 can induce the amyloid aggregation of its paralogs, p63 and p73, from the droplet state advances our understanding of the molecular underpinnings of oncogenesis. The results are summarized in a scheme in Figure 7. This phenomenon highlights the dual nature of p53 as both a guardian against and a facilitator of tumorigenesis, depending on its conformational state.

**Figure 7.**
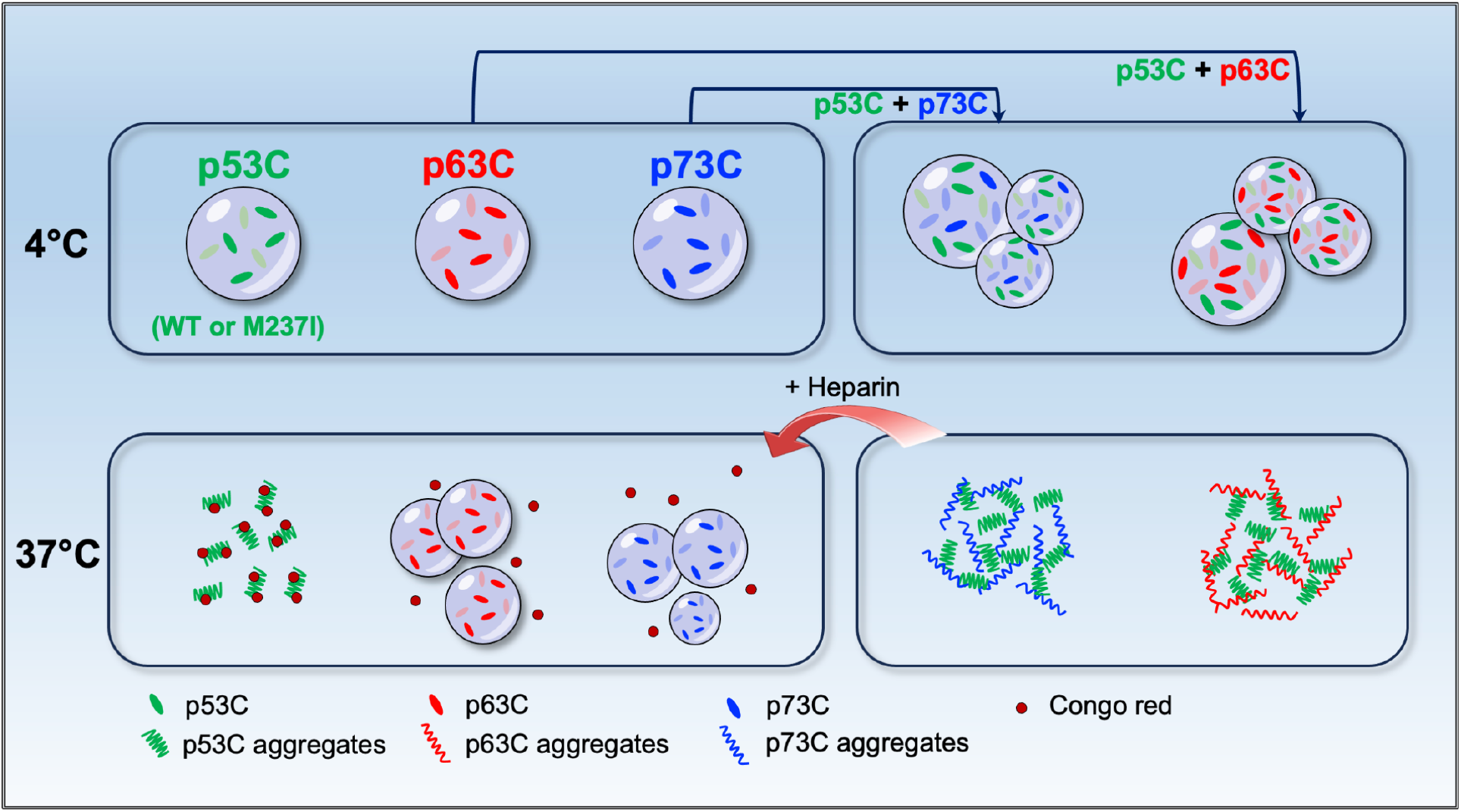
Scheme showing the effects of p53C and M237I converting p63C and p73C droplets into amyloid aggregates at physiologically relevant temperatures.

The capacity of both mutant and WT p53 to induce a shift in p63 and p73 from a liquid to an amyloid-like solid state implies a potential universality in the aggregation mechanisms of these proteins. This insight is paramount as it suggests that the oncogenic potential of p53 may not be solely relegated to its mutant forms. Instead, it positions the phase behavior of p53 as a central factor in the dysregulation of tumor suppressor functions^11, 31, 37^.

In non-cancerous cells, p53 is expressed at low concentration with half-time of minutes. There is an increase in p53 concentration only under stress; under some circumstances, the protein eventually undergoes phase separation into a condensate, where it exerts its function^32^. The observed heterotypic phase transition involving p53 and the paralogs p63 and p73 provides a new perspective on the gain of function (GoF) associated with mutant p53. It is tempting to associate the oncogenic properties of mutant p53 with its propensity to alter the phase state of other proteins rather than solely from direct genetic alterations. These findings align with emerging models of cancer that emphasize the significance of protein-protein interactions and the tumor microenvironment in disease progression. It is tempting to suggest that the aberrant condensates of mutant p53 may act as a scaffold on p63 and p73 and other transcription factors.

Recent research has unveiled that heterotypic condensates composed of prion protein and α-synuclein can transition from a liquid to a solid state, a transformation that is particularly modulated by RNA, resulting in the formation of heterotypic amyloids^38^. The same research team has observed that tau protein and PrP droplets mature as they age, culminating in a solid-like state characterized by amorphous and amyloid-like co-aggregates^39^. These findings highlight the complex interactions between proteins implicated in neurodegenerative diseases. Our study introduces a novel perspective by demonstrating that heterotypic interactions between p53 and its paralogs, p63 and p73, can lead to amyloid aggregation. This marks the first instance of such an effect in cancer-associated proteins, broadening our understanding of protein phase transitions in oncology. Evidence indicates that p53 amyloid aggregates can entrap p63 and p73 in both in vitro and in vivo settings^7, 21^, but the underlying mechanisms of this sequestration remained obscure. Our findings elucidate that p53 aggregates directly engage with biomolecular condensates of p63 and p73 (Figure 6), providing a mechanistic understanding of this interaction.

Moreover, the prion-like conversion of p63 and p73 by mutant p53 broadens the scope of potential therapeutic targets within the p53 signaling cascade^11, 12, 40–44^. The selectivity of these interactions provides a unique opportunity to design inhibitors that specifically disrupt the aberrant phase transitions while preserving the normal functions of these proteins. Numerous studies have identified p53 aggregation as a promising focus for anti-cancer therapies^11, 12, 19, 41, 45–48^. The recent discovery that antineoplastic drugs are preferentially localized within specific protein condensates in the nuclei of tumor cells^42, 43^ underscores the significance of our findings regarding the conversion of non-amyloidogenic tumor suppressors p63 and p73 by mutant p53. This observation suggests that drugs designed to target p53 aggregation might selectively accumulate in the condensates that contain p53 and its related proteins, enhancing the specificity and efficacy of therapeutic interventions.

Our experiments with heparin serve as a proof of principle, demonstrating that the seeding-induced transition of p53 to an aggregated state, affecting its paralogs p63 and p73, can be halted (as shown in Figure 5). Heparin not only arrests the transition of p53 into the amyloid state but also inhibits the seeding of p53 on the liquid droplets of p63 and p73 (Figures 5 and 7). This striking observation presents potential therapeutic avenues to be explored using similar polyanions, which could include specific RNA or DNA sequences and certain small molecules.

While our research has advanced the understanding of p53 phase transitions, certain aspects warrant further investigation. The complexity of the nuclear environment, with its abundance of nucleic acids, appears to influence p53 phase separation and aggregation, as referenced in several studies^11, 49–54^. Variables such as the RNA-to-protein ratio have been shown to modulate the phase transition of mutant p53^31, 51, 53^, and this modulation extends to its paralogs, as demonstrated in our heparin experiments. Additionally, the broader spectrum of environmental and genetic factors that may impact this process within the cancer cell milieu is ripe for detailed investigation. The role of post-translational modifications on the phase behavior of these proteins also presents a promising avenue for research. Pursuing in vivo studies will be crucial to expand upon our findings and deepen the understanding of how targeting phase transitions in the p53 protein family may hold therapeutic value.

In conclusion, our study contributes a significant piece to the puzzle of p53 in cancer biology. By delineating the conditions that precipitate the phase transition and amyloid aggregation of p63 and p73, we have identified a potential vulnerability in cancer cells that could be exploited in developing novel anticancer therapies. Future work focusing on developing small molecules or biologics that can prevent or reverse the amyloid aggregation of p53 paralogs holds the promise of advancing cancer treatment strategies. The path ahead is challenging but filled with the potential for significant breakthroughs in the fight against cancer.

## Methods

### Protein Expression and Purification

The genes encoding p53C WT and M237I, p63C, and p73C proteins were subcloned into pET15-b (Addgene, #24866), pET28-a, and pNIC-CTHF vectors (Addgene, #39077), respectively, and were expressed in *Escherichia coli* BL21-CodonPlus. Cultures were grown in Luria-Bertani medium and incubated at 37°C until the optical density (OD_600_) of 0.8 was reached. Protein expression was induced with 1 mM isopropyl-β-D-thiogalactopyranoside (IPTG) overnight at 25°C under agitation. Cells were harvested by centrifugation, resuspended in buffer (pH 7.4) containing 50 mM Tris-Cl, 500 mM NaCl, 5 mM dithiothreitol (DTT), and 1 mM phenyl-methyl-sulfoxide (PMSF) protease and disrupted by sonication. Insoluble proteins and cell debris were separated by centrifugation. Highly homogenous samples of p53C WT, p53C M237I, and p63C were obtained with the following chromatography steps: soluble fraction of each protein was loaded onto a HisTrap FF nickel affinity column (Cytiva,) and elution was performed with a linear increasing imidazole gradient. Cleavage of the fusion tag was achieved by thrombin digestion [1:1000 w/w] at 4°C overnight. The product was diluted and loaded onto a Heparin Sepharose FF affinity column. The p73C purification was described elsewhere^21^. Briefly, a soluble fraction of p73C was loaded onto the HisTrap FF nickel affinity column (Cytiva), digested with Tobacco Etch Virus (TEV) protease [1:50 w/w], reapplied in a HisTrap FF column (Cityva) and the non-bound was collected. All final samples were dialyzed against 20 mM Tris-Cl (pH 7.4), 150 mM NaCl, 5 mM DTT, 5 µM ZnCl_2_, 5% glycerol and stored at −80°C. Before each experiment, all proteins were thawed on ice, centrifuged at 14,000 g 4°C for 15 minutes, and passed through a 100 KDa microcon centrifugal filter to exclude residual aggregation. Protein final concentration for each experiment was measured using absorbance at 280 nm and extinction coefficient of 18,040 M^-1^cm^-1^ for p53C WT and M237I, 15,930 M^-1^cm^-1^ for p63C, and 18,91 M^-1^cm^-1^ for p73C.

### Protein labeling

For some LLPS microscope experiments, p63C and p73C were labeled with NHS-Rhodamine (Thermo Scientific) and p53C WT and M237I with Alexa FluorTM 488 (Thermo Scientific) fluorescent dyes using the manufacturer’s specifications. Each protein was dialyzed against 0.1 M sodium bicarbonate pH 8.5 and incubated on ice for 2 h with a five-fold molar excess of NHS-Rhodamine or one vial of Alexa FluorTM 488. The dye excess was separated using a HiTrap Sephadex G-25 Desalting column (Cytiva), and proteins were eluted in 20 mM Tris-Cl (pH 7.4), 150 mM NaCl, 5 mM DTT, 5 µM ZnCl_2_, 5% glycerol. Protein concentration and labeling efficiency were calculated as described in each manufacturer’s user guide.

### Differential interference contrast microscopy (DIC) and Fluorescence microscopy

DIC and fluorescence images were captured using a Zeiss LSM 710 confocal laser scanning microscope equipped with a Plan-Apochromat 63x/1.4 objective lens. Supernatants were combined with a pre-chilled 50% PEG-4000 stock solution at 4°C, resulting in a final PEG concentration of 15% for all experiments unless otherwise specified. Samples (20 μL) in 20 mM Tris-Cl (pH 7.4), 150 mM NaCl, 5 mM DTT, and 5 µM ZnCl_2_ were loaded onto coverslips, and imaging was conducted at room temperature. For concentration-dependent experiments involving p63C and p73C (Figure 1a), PEG-4000 was added to protein solutions at concentrations ranging from 10 to 100 µM (at 4°C), and images were captured using DIC microscopy. Temperature-dependent experiments involved incubating proteins at concentrations of 60 μM across temperatures of 4, 25, or 37°C for different periods, including 3 minutes (Figure 1c), and extending to 30, 90, or 270 minutes (Figure 1d). To perform the Congo Red (CR) binding assay (Figure 2 a-d), proteins at concentrations of 60 μM were initially incubated at 4 or 37°C for 3 min before the addition of CR (10 µM). After the CR addition, PEG-4000 was added to the sample, which was then proceeded to analysis.

The influence of wild-type p53C on p63C and p73C aggregation dynamics (Figure 3) involved mixing NHS-Rhodamine labeled p63C or p73C (40 μM) with wild-type p53C (40 μM) and PEG-4000. Additionally, aggregation of p63C and p73C induced by wild-type p53C and M237I in the presence of PEG (Figure 4a-d) was investigated, where NHS-Rhodamine labeled p63C or p73C (40 μM) was combined with Alexa-Fluor488-labeled wild type p53C or M237I (40 μM) and PEG-4000 for imaging at 4 and 37 °C. For seeding of p53C and heparin effects on p63C droplets conversion to aggregates (Figure 5f-g), NHS-Rhodamine labeled p63C (40 μM) was combined with Alexa-Fluor488-labeled wild-type p53C or M237I (40 μM) at 37 °C for 3 minutes before the addition (or not) of heparin (10 µM). Then, PEG-4000 was introduced, and the sample was imaged. Image processing was performed using Fiji, a distribution package of ImageJ software.

### Thioflavin T and light scattering measurements

Light scattering (LS) and Thioflavin T experiments were performed in an ISSK2 spectrofluorometer (ISS Inc). To investigate the influence of temperature in LLPS/aggregation, wt p53C, M237I, p63C, and p73C were prepared in a buffer containing 20 mM Tris-Cl, (pH 7.4), 150 mM NaCl, 5 mM DTT, 5 µM ZnCl_2_, and 5% glycerol. The proteins were then subjected to pre-incubation at concentrations of 60 μM, with or without the addition of 15% (w/v) PEG-4000, for a duration of 3 minutes across different temperatures: 4°C, 25°C or 37°C. Samples were excited at 320 nm, and emission was recorded from 300 to 340 nm. Experiments were performed with a 5 nm slit width and 90% of the iris closed. Data were expressed as the area under the LS curve. Samples in the same condition were incubated with Thioflavin T at 10 μM final concentration, excited at 450 nm, and emission was recorded from 460 to 500 nm. Thioflavin T emission was expressed as intensity at 470 nm.

Light scattering and Thioflavin T co-aggregation experiments were conducted as described above with 40 μM of p53C or M237I and 40 μM p63C in the same buffer in the presence or absence of 15% (w/v) PEG-4000. Mixtures were pre-incubated at 4°C and 37°C for 3 minutes before each measurement. Heparin at a final concentration of 10 μM was used to investigate its modulatory properties during the aggregation process. For seeding experiments, a stock solution of p53C or M237I at 60 μM final concentration was submitted to aggregation at 37°C for 3 minutes. Then, it was diluted to a final concentration of 1.5 μM and mixed with 60 μM of p63C in the presence of 15% (w/v) PEG-4000. Light scattering and Thioflavin T binding were investigated as described before.

### Turbidity measurements

Turbidity assays were performed in a Varioskan Lux 3020-81205 Microplate Reader (Thermo Scientific). P53C, M237I, p63C, and p73C dissolved in a solution of 20 mM Tris-Cl (pH 7.4), 150 mM NaCl, 5 mM DTT and 5 µM ZnCl_2_, with15% (w/v) PEG-4000 at concentrations ranging from 10 to 100 µM were transferred to a 96 well optical black/clear bottom plate (Thermo Scientific) and pre-incubated for 3 min at 4°C, 25°C or 37°C. Turbidity was measured as absorbance at 600 nm.

### Cell Culture

Human non-small lung carcinoma H1299 cells were obtained from The Rio de Janeiro Cell Bank (RJCB) and cultured in RPMI-1640 medium supplemented with 10% fetal bovine serum. Cells were maintained at 37°C in an atmosphere containing 5% CO2.

### Cell Transfection

Transfection experiments were performed following the Lipofectamine 2000 reagent (Invitrogen) protocol provided by the manufacturer. The day before transfection, 5×104 cells were cultured in a 24-well plate containing a glass coverslip. After reaching 70% of confluency, H1299 cells were transfected with ten µg of pEGFP-N1 vector (Clontech), containing a C-terminus EGFP-tagged sequence of the full-length M237I p53 protein (Addgene, #11770), and 4 µL of Lipofectamine 2000 reagent. After 24 hours since transfection, the immunofluorescence protocol was performed.

### Immunofluorescence

The H1299 cells were washed thrice with PBS and fixed with paraformaldehyde 4% for 20 minutes at room temperature. Then, cells were washed with PBS and permeabilized with Triton X-100 (0.5%) for 15 minutes. After another wash step with PBS, cells were incubated with a 3% bovine serum albumin (BSA) (Sigma) solution for 1 hour at room temperature. Cells were labeled with monoclonal anti-p73 antibodies in three different dilutions: 1:50, 1:100, and 1:200 overnight at 4°C. After that, cells were washed with PBS and incubated with a 1:1,000 dilution of Alexa Fluor 568 antibody for 1 hour at room temperature in the dark. Finally, cells were washed with PBS and fixed with Vectashield Antifade Mounting Medium with DAPI. Images were acquired with an ELYRA LSM 710 confocal laser scanning microscope (Carl Zeiss, Inc.).

## Acknowledgments

Our laboratory is supported by grants from the National Council for Scientific and Technological Development (CNPq awards and the INCT program, grant no. 465395/2014-7 and 408046/ 2021-0 to J.L.S. and grant no. 313137/2021-8 to G.A.P.d.O.) and the Carlos Chagas Filho Foundation for research support in the state of Rio de Janeiro (FAPERJ) grant no. 210.008/2018 and 202840/2018 to J.L.S., grants E-26/201.296/2021 and E26/210.294/2022 to G.A.P.d.O. and grants E-26/200.582/2022, E-26/210.346/2022 to M.A.M and E-26/201.325/2021 to T.C.R.G.V.

## Authors contributions

All authors listed have made a substantial, direct, and intellectual contribution to the work and approved it for publication. E.C.P., G.C.A., G.S.S., F.P.A., M.F.M., and M.A.M. performed experiments; E.C.P. and G.C.A. writing the original draft; T.C.R.G.V., G.A.P.d.O., and J.L.S. conceptualization, data curation, funding acquisition, project administration, editing the manuscript and figures, and supervising the research.

## Notes

### Competing Interest Statement

The authors have declared no competing interest.

