## Supplementary Figures for "Oncogenic Phase Transitions: How Mutant p53 Drives Amyloid Formation in p63 and p73 Liquid Droplets"

*Equally first authors

### Corresponding authors

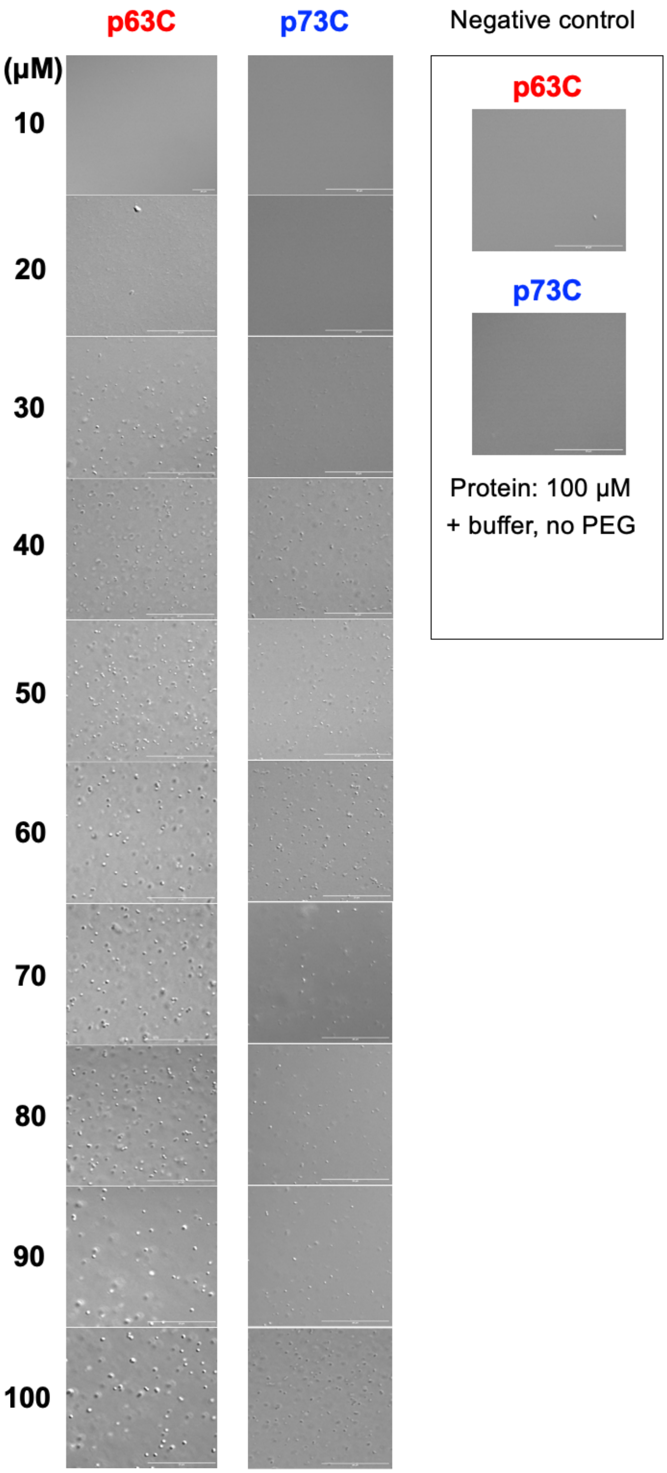


**Supplementary figure 1.** Representative DIC images at increasing p63C and p73C concentrations (ranging from 10 to 100 μM). Control experiments show the absence of p63C and p73C droplets when PEG is withdrawn from the solution. Scale bar: 20 μm.

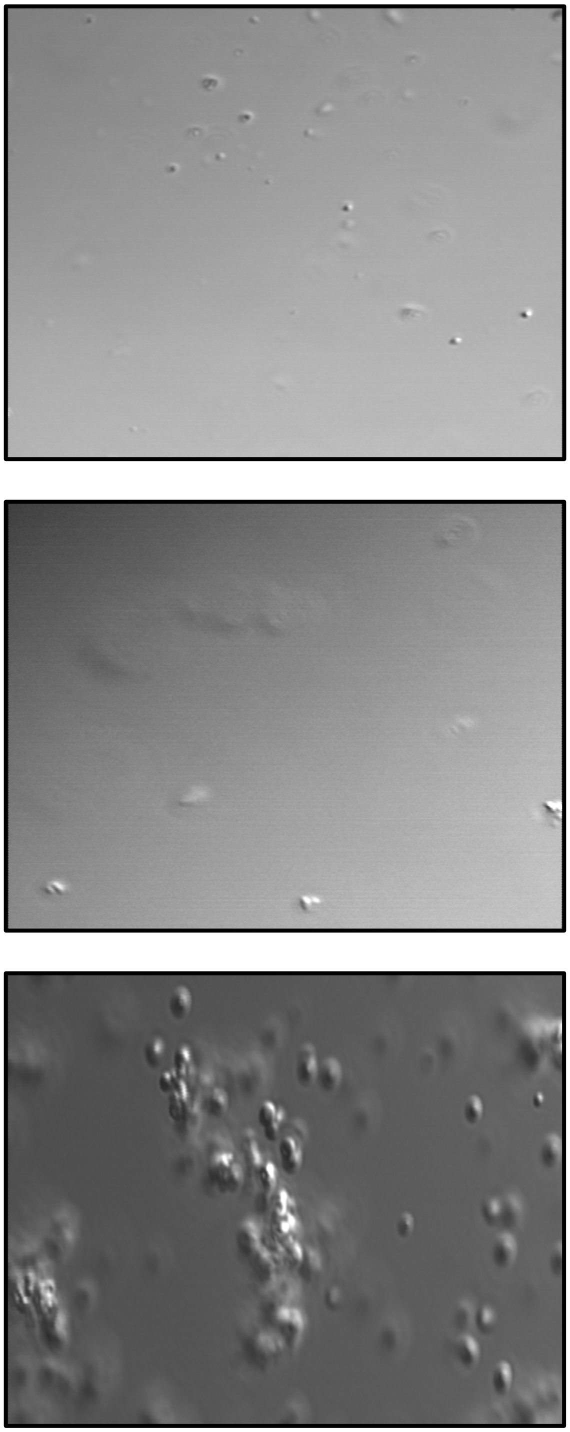

**Supplementary figure 2.** Videos obtained from the bright-field channel at early (after 30 minutes, upper), intermediate (after 90 minutes, central), and later (after 270 minutes) stages of p53C. (See methods for details).

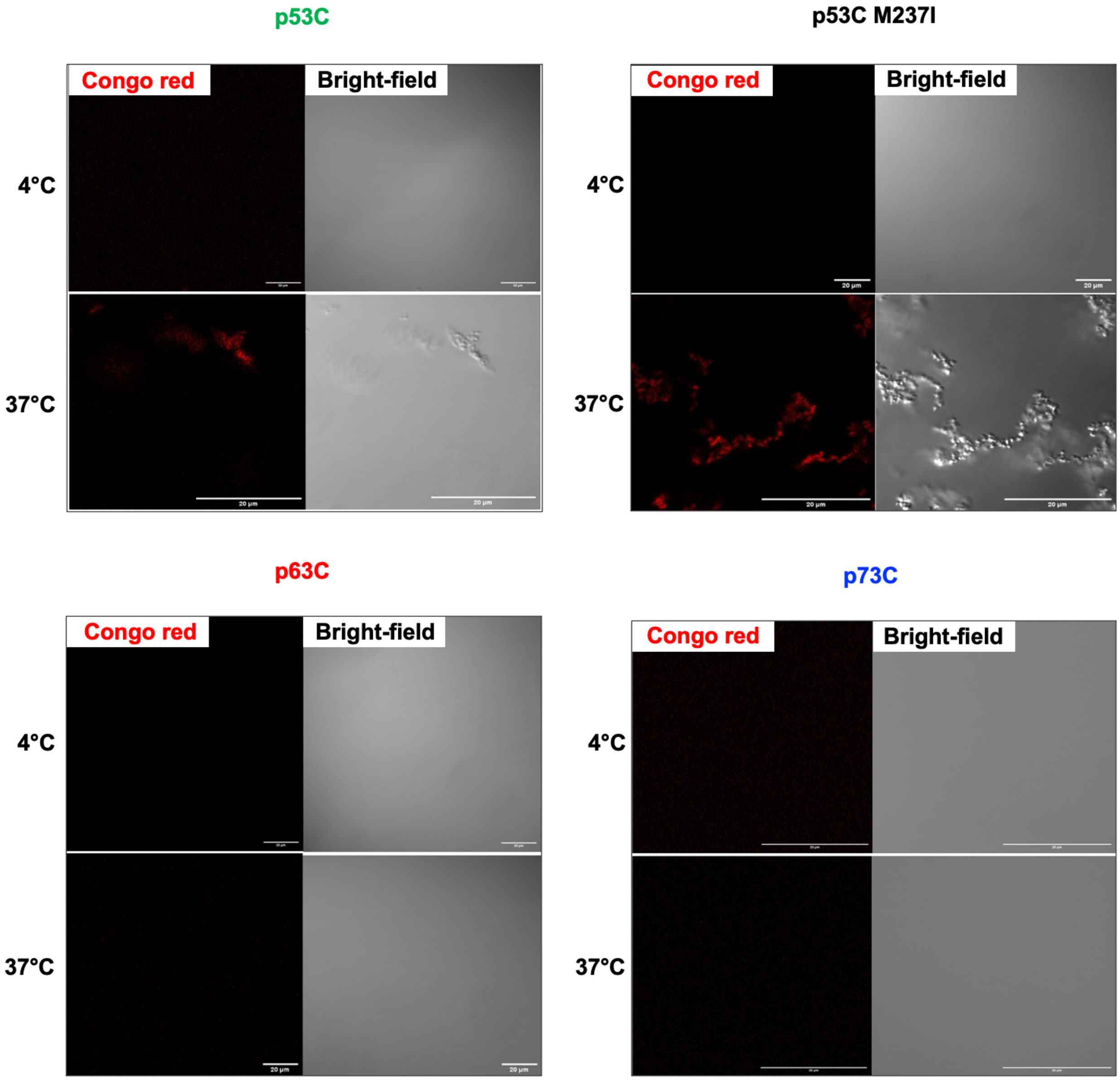


**Supplementary figure 3.** Control experiments of Congo red binding to the p53C family members in the absence of PEG. Congo red (CR) fluorescent and bright-field channels of p53C, M237I, p63C, and p73C at 4 and 37°C. Scale bar: 20 µm.

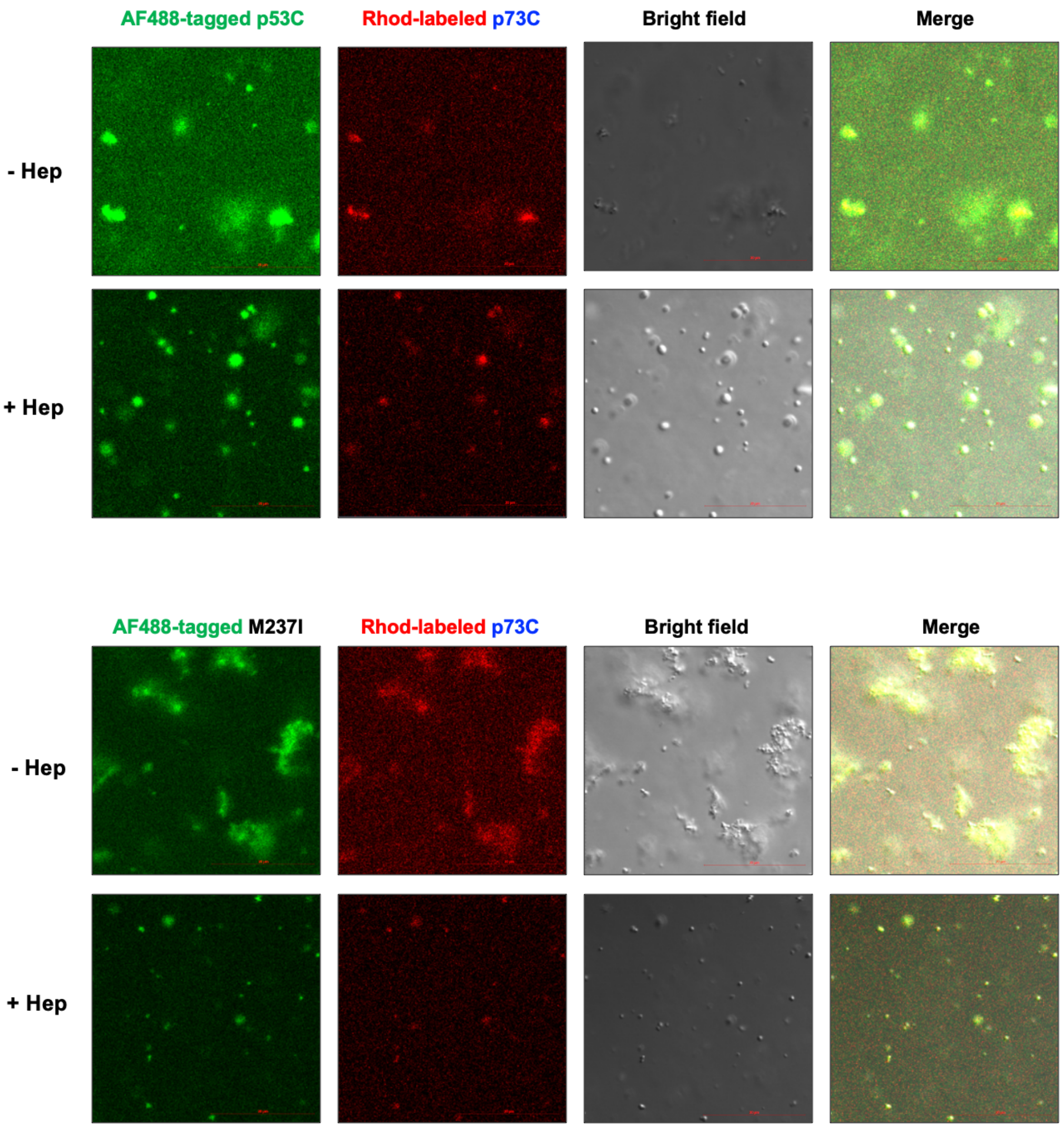


**Supplementary figure 4.** Collection of DIC images showing the AF488-tagged p53C (upper) or M237I (lower), the Rhodamine-labeled p73C, the bright-field, and merged channels in the absence (-) or presence (+) of heparin. Scale bar: 20 μm.

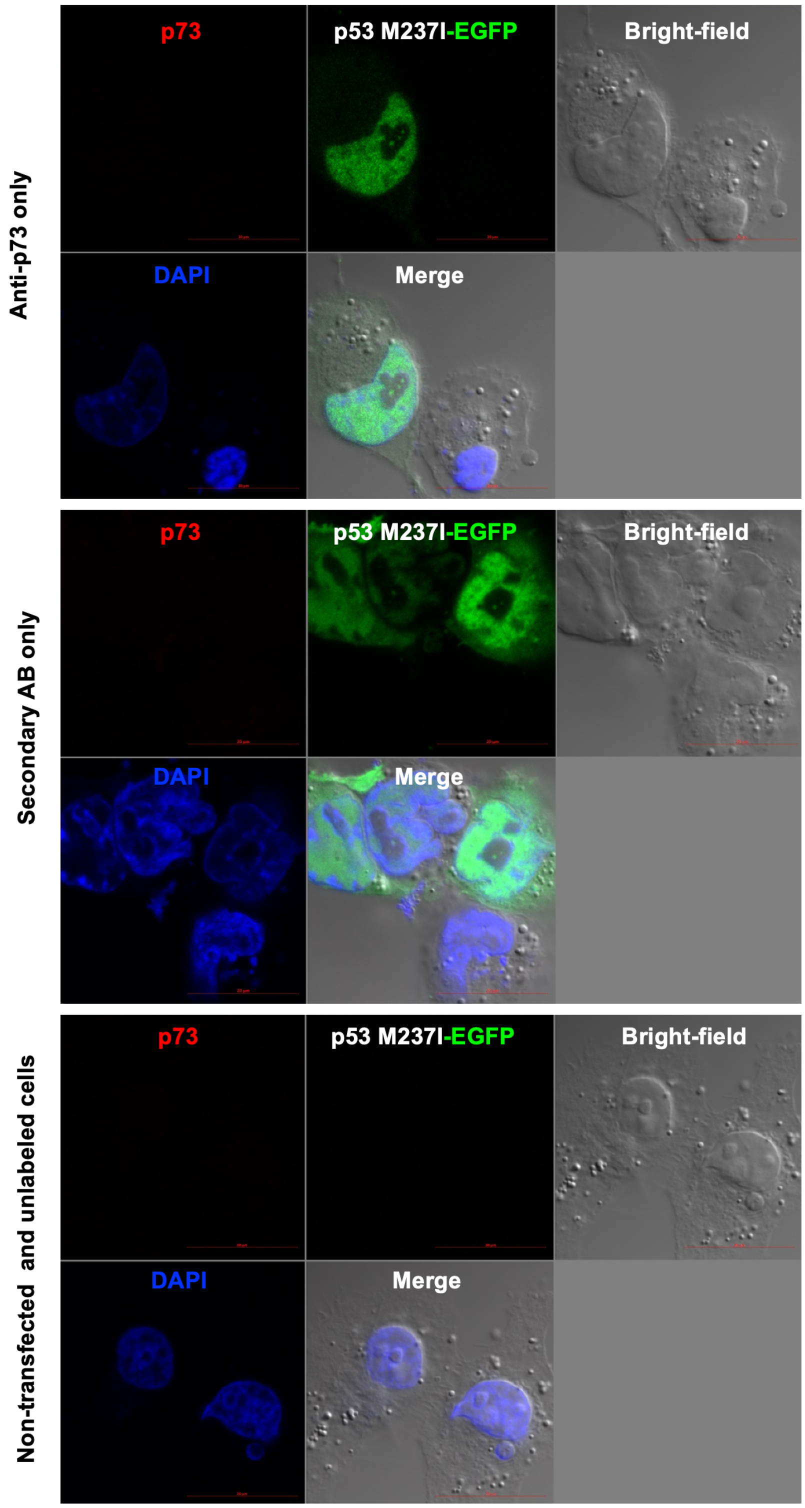


**Supplementary figure 5.** Immunofluorescence control experiments. Collection of microscopy images showing regions of interest after performing immunostaining with anti-p73 only (top), secondary antibody (AB) only (middle), and non-transfected and unlabeled cells (bottom).
